# redkmer: an assembly-free pipeline for the identification of abundant and specific X-chromosome target sequences for X-shredding by CRISPR endonucleases

**DOI:** 10.1101/189431

**Authors:** Philippos Aris Papathanos, Nikolai Windbichler

**Affiliations:** Department of Experimental Medicine, University of Perugia; Department of Life Sciences, Imperial College London; Corresponding author

## Abstract

CRISPR-based synthetic sex ratio distorters, that operate by shredding the X-chromosome during male meiosis, are promising tools for the area-wide control of harmful insect pest or disease vector species. However, the selection of gRNA targets, in the form of high-copy sequence repeats on the X chromosome of a given species, is difficult since such repeats are not accurately resolved in genome assemblies and can’t be assigned to chromosomes with confidence. We have therefore developed the redkmer computational pipeline, designed to identify short and highly-abundant sequence elements occurring uniquely on the X-chromosome. Redkmer was designed to use as input exclusively raw WGS data from males and females. We tested redkmer with suitable short and long read WGS data of *An. gambiae*, the major vector of human malaria, in which the X-shredding paradigm was originally developed. Redkmer establishes long reads as chromosomal proxies with excellent correlation to the genome assembly and uses them to rank X-candidate kmers for their level of X-specificity and abundance. Redkmer identified a high-confidence set of 25-mers, many of which belong to previously known X-chromosome specific repeats of *An. gambiae*, including the ribosomal gene array and the selfish genetics elements harbored within it. WGS data from a control strain in which these repeats are also present on the Y chromosome confirmed the elimination of these kmers in the filtering steps. Finally, we show that redkmer output can be linked directly to gRNA selection and can also inform gRNA off-target prediction. The redkmer pipeline is designed to enable the generation of synthetic sex ratio distorters for the control of harmful insect species of medical or agricultural importance. It proceeds from WGS input data to deliver candidate X-specific CRISPR gRNA candidate target sequences. In addition the output of redkmer, including the prediction of chromosomal origin of single-molecule long reads and chromosome specific kmers, could also be used for the characterization of other biologically relevant sex chromosome sequences, a task that is frequently hampered by the repetitiveness of sex chromosome sequence content.

## Introduction

We have previously shown in the malaria mosquito *An. gambiae* that sex ratio distorters can be rationally engineered using endonuclease-mediated X-chromosome shredding during spermatogenesis [3, 4]. We also have shown that sex ratio distortion is a useful phenotype to engineer in pest or vector insects, as it can lead to population suppression in a manner that, as predicted [19], is more efficient than the classical Sterile Insect Technique (SIT). Furthermore, X-shredders are attractive options for vector control because physically linking these to a Y chromosome can provide both the allele itself and the entire Y chromosome which harbors it a competitive advantage in inheritance against the X [7]. Through this benefit such an engineered Y chromosome can display genetic drive increasing in frequency within a population rapidly starting from low frequencies, biasing the sex ratio towards males as it increases. As a result, the reproductive potential of the population diminishes for the lack of females. Because a large number sites targeted simultaneously through X-shredding, the development of resistance mechanisms is significantly impaired, unlike other strategies involving endonucleases for population suppression [8, 20].

Significant interest has developed in the genetic control community to engineer such synthetic sex ratio distorters in additional insect pests or disease vectors, since X-shredding exploits and manipulates the near universal significance of paternal chromosome inheritance on the sex of an individual. There are five essential requirements required to do so: (1) an XY male karyotype, (2) genetic transformation, (3) regulatory elements (promoters) that can drive expression of the X-shredding nuclease during spermatogenesis, (4) an endonuclease platform such as the CRISPR/Cas9 system that can be directed against X-chromosome specific sequences and finally (5) the existence of sequences on the X chromosome that are both specific and abundant to it.

In our previous work in the malaria mosquito, we were able to build an X-shredder because this mosquito’s rDNA genes are exclusively located on the X-chromosome. However, this arrangement is exceptional and the vast majority of insects do not to share it. Furthermore, knowledge of naturally occurring multi-copy X-specific sequences, e.g. X-specific satellite DNA, is limited because such repetitive DNA sequences are *ipso facto* excluded from genome assemblies and because few studies deal with such elements particularly in non-model organisms. Indeed, even after 15 years since the publication of the first genome assembly of *An. gambiae* [9], the rDNA cluster is still not correctly represented in the current genome assembly. Knowledge of the rDNAs’ specificity to the X chromosome in *An. gambiae* came from studies of mosquito population genetics using cytology. To address this limitation we have developed a bioinformatic pipeline called redkmer, for repeat extraction and detection based on k-mers, that by utilizing long and short read sequencing technology, is able to identify highly abundant, X-specific sequences in the absence of a genome assembly.

## Methods

### Data requirements

Redkmer requires as input whole genome sequencing (WGS) data based on long single-molecule (e.g. PacBio) and short (e.g. Illumina 100bp) reads. For the long reads, WGS must be performed using male-only or mixed sex samples and reads must be self-error corrected, for example using Canu [11], and provided in fasta format. For the short read libraries, data must be generated from both male and female samples independently and provided in fastq format as a single file (paired end reads can me merged into one file for each sex). A fasta file with the mitochondrial reference genome is also required to remove short reads derived from it. Redkmer is optimized to run with data sets that achieve at least a 10x genome coverage of the long read library and a 20x genome coverage of each short read library.

### Software and hardware requirements

Redkmer has been implemented to run on UNIX HPC systems scheduled by SGE or PBS. All steps were run using the cx1 general purpose cluster at the Imperial College High Performance Computing Service. All alignment steps allow splitting of the long read input data via the [NODES] parameter for parallel execution, and have been run on standard 12 and 24 core nodes with 32GB of memory. Processing steps require up to 120GB of memory depending on the size of the input libraries. The following third party modules are required and loaded by redkmer: bowtie2 [14], bowtie1 [13], fastQC [18], jellyfish [16], samtools [15], BLAST [1] and the R environment for plotting which utilizes ggplot2 and data table modules.

### Pipeline overview

Redkmer is designed to identify short 25bp sequences (kmers) occurring abundantly and specifically on X-chromosomes of species with XY male karyotypes. The pipeline is designed around three basic principles for target site identification and selection: (1) X chromosome kmers are first identified by assessing differential representation in female vs male WGS data, also known as chromosome quotient (CQ) [6, 5]. (2) kmers that occur on other chromosomes are subsequently eliminated and (3) those that are the most abundant within those displaying X-chromosome specificity are selected. In the first phase of the pipeline, redkmer uses unassembled error-corrected reads from single-molecule WGS (called PacBio from here for simplicity) as chromosomal proxies, by mapping to them short reads from separate female and male WGS libraries (called Illumina from here for simplicity). The ratio of mapping female and male Illumina reads, also known as the chromosome quotient (CQ), is used to predict chromosomal origin, where reads originating from the X chromosome typically display CQ of 2. Chromosomal bins are thus populated by PacBio reads, which are assigned to one of four bins (autosomal, X, Y and Genome-Amplification - GA) depending on their CQ. Because CQ is calculated through the mapping of many Illumina reads to a relatively longer sequence, the confidence in the predicted chromosomal origin of PacBio reads is higher compared to CQ calculated over the span of only the target site. Furthermore, on account of PacBio read length, redkmer can distinguish between reads derived from the X chromosome and those that are autosomal in origin, but which harbor sequences homologous to those on the X chromosome, which depends on the relative length of the match and the length of the PacBio read. In the second phase, kmers generated from the Illumina libraries are mapped to PacBio reads of each chromosomal bin separately. Kmers mapping above a defined threshold to non-X derived Pacbio reads compared to X-reads are tagged as unsuitable for X-shredding. Redkmer also generates data regarding off-targeting potential of candidate X-specific kmers by searching for degenerate target sites in non-X chromosome derived reads. The output of the pipeline, a list of suitable kmers along with their specificity and abundance profiles, can then be used for the purpose of to building and testing of X-shredder constructs, by feeding selected kmers into external computational tools that predict their suitability for RNA-guided nuclease platforms, for example for CRIPR/Cas9.

### Implementation and module functionality

Redkmer is freely available in Github (<u>github.com/genome-traffic/redkmer-hpc</u>)

under the GNU General Public License Version 3, 29 June 2007. Redkmer implements ten core modules (Figure 1) and is designed to run on a high performance computing platform. All configurable parameters and run settings are controlled from the file *redkmer.cfg* that sets redkmer behavior. Redkmer first runs quality control and filtering of the input data. Reads from Illumina libraries are mapped to the mitochondrial genome using bowtie2 [13] and aligned reads are removed from each library. This step is required, because mitochondrial sequences can behave similarly to X chromosomal sequences when redkmer calculates CQ, i.e. having a higher coverage in females compared to males (e.g. in Drosophila whole body WGS). The quality of the filtered Illumina reads is then checked with FastQC [18] and reports are generated for the user. PacBio reads are then filtered for read length removing those of insufficient length to reliably predict chromosomal origin (Module 1). Redkmer then assigns the PacBio reads to chromosomal bins, by separately mapping Illumina reads from the male and female libraries to the filtered PacBio reads using bowtie [14], allowing no mismatches throughout the length of the alignment, reporting all possible alignments and normalizing the number of mapping reads for the library sizes (Modules 2-3). The initial kmer sets are generated from the female and male Illumina WGS libraries separately using jellyfish [16], with which redkmer counts kmer abundance in both libraries and calculates their ratio of counts (effectively kmer-CQ) normalized by library size (Modules 4-5). All kmers are then mapped to reads in each chromosomal bin using bowtie [14]. The number of hits to each chromosomal bin is counted and through this redkmer derives the X-specific index (XSI) as the ratio of X-chromosome over non-X hits. Therefore, redkmer selection does not exclude kmers with perfect off-targets (100% matches of the kmer to the non-X bins) using an arbitrary cutoff, but instead uses the proportional cutoff that accounts for the total number of kmer hits to non-X reads (Modules 6-7). Kmers with a XSI that pass a selected threshold (e.g. 0.99 - less that 1% of hits tolerated to non-X reads) are then remapped to all non-X long reads, this time allowing 20% mismatches over the length of the alignment to identify “degenerate” off-targets (Modules 8-9). Redkmer finally processes the outputs of all modules and generates fasta and tab-delimited list of candidate kmers for X-shredding (Module 10). There are a number of supplementary modules that have been built to evaluate redkmer output for example against an available genome assembly, or set of reference genes located in the supplementary modules folder.

**Figure 1:**
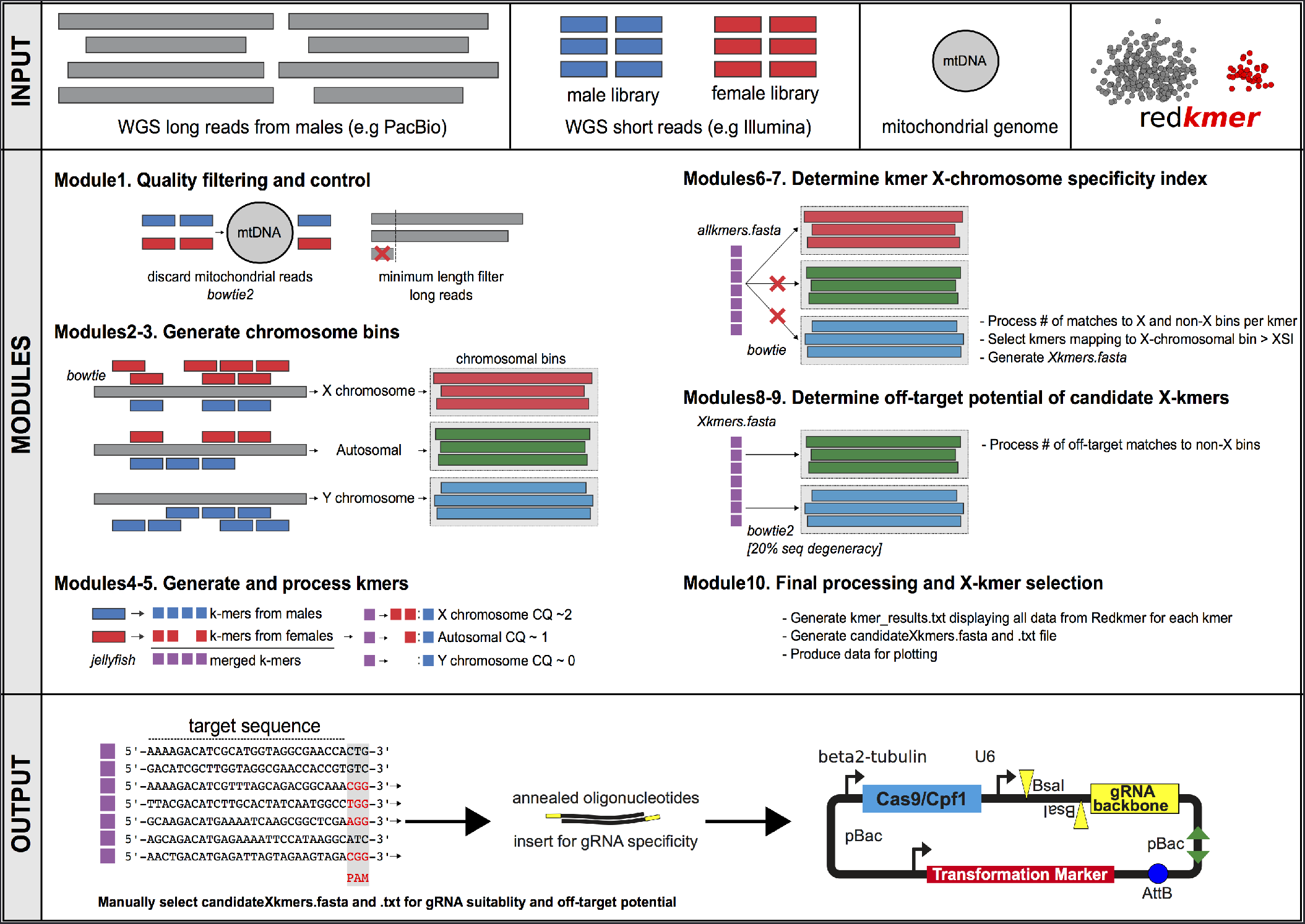
Schematic overview of the redkmer pipeline showing required input data, module main functions and potential downstream activities

## Results

### Redkmer target selection in *An. gambiae*

To test redkmer target selection we ran it using WGS data from the Pimperena strain of *An. gambiae*, which are publicly available at the Sequence Read Archive (SRS667972, SRR1509742 and SRR1508169). Annotated genomic data, for example the mitochondrial genome and the genome assembly AgamP4 were based on the PEST strain and retrieved from Vectorbase. We selected *An. gambiae* for redkmer evaluation because the rDNA gene cluster and the repetitive elements residing within it are already known to be both X-specific and abundant and have been experimentally shown to be well suited for X-shredding using both homing endonucleases as well as CRISPR based endonucleases. In addition, separate male and female Illumina WGS data are also available for the Asembo1 strain of *An. gambiae* (SRR1504990 and SRR1504983), which was used here as a control, as this strain harbors the ribosomal gene cluster on both sex chromosomes, likely as a result of a rare X-Y recombination event [21].

### Pacbio reads as chromosome proxies

Of the initial ∼4.3million error-corrected input PacBio reads, redkmer retained ∼2 million reads that passed the minimum length cutoff(default setting of 2kbp), resulting in a total of 7.4x10^9^ sequenced nucleotides (∼25x coverage assuming a 300Mbp genome). The filtered PacBio reads were assigned to one of the four chromosome bins based on read CQ from Modules 2-3 (Figure 2A). PacBio read CQ and coverage (measured in length normalized sum of Illumina reads from males and females mapping to each PacBio read - LSum) indicated that both autosomes and the sex chromosomes harbor numerous repetitive elements (Figures 2B). Importantly, the high density of PacBio reads with high LSum and with a CQ close to 2 indicated an abundance of repeats on the X chromosome and not shared with other chromosomes (Figure 2B). A significant co-occurrence of X-linked repeats on the autosomes or the Y chromosome would result in significant shift of the CQ towards 1, which was not apparent for the repeat-containing PacBio reads of the X-bin (Figure 2B). Overall, PacBio reads from the Y chromosome bin displayed the highest level of repetitive DNA content, followed by X chromosomes and then the autosomes (Figure 2C), consistent with published data [5]. Reads belonging to the GA-bin had the lowest LSum values, consistent with the explanation that reads with CQ significantly higher than 2 would be expected either to result from sample specific sequencing artifacts or bacterial contamination (Figure 2C). Additional redkmer generated plots, providing data on basic statistics of the input PacBio reads library including the distribution of CQ, LSum, Sum (sum of mapping reads from males and females prior to normalization) and read length can be found in Supplementary Figures 1-8.

**Figure 2:**
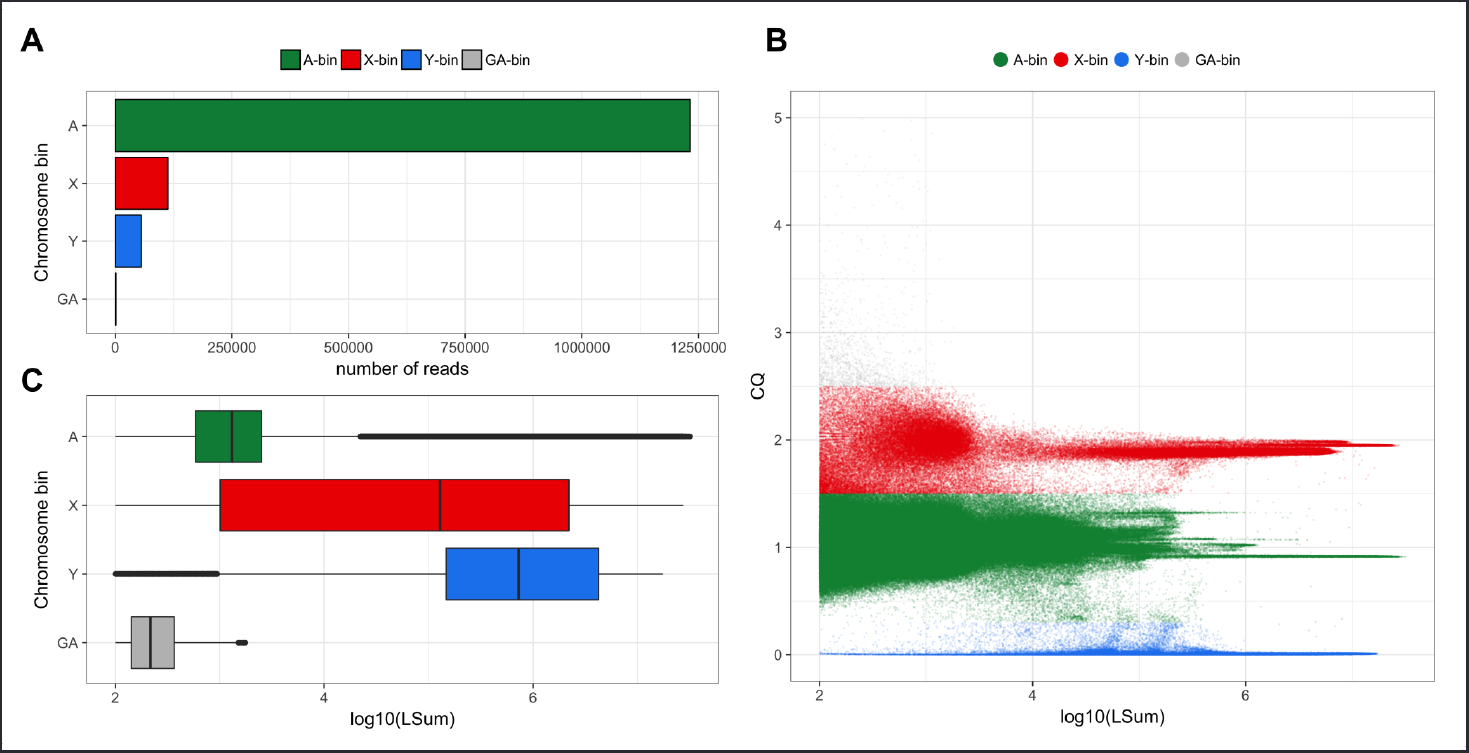
Long read assignment into chromosomal bins using the *A. gambiae* Pimperena strain. **A)** Number of PacBio reads assigned to each chromosome bin. B) PacBio read CQ (ratio of female/male data) over log10 of LSum (total number of mapping reads from male and female illumina data) showing chromosomal bins and chromosome repetitiveness. C) Box-plots of PacBio coverage/repetitiveness for each chromosomal bin.

To evaluate CQ-based prediction of chromosome origin, we mapped PacBio reads from each bin by BLAST [1] to the latest *An. gambiae* genome assembly (Table 1, Figure 3) in a non-exclusive manner reporting all matches of 2kb or above. Hits to the 42.39 Mbp long “unknown chromosome”, which contains all repeat-rich scaffolds that have not been anchored to chromosomes by physical mapping, accounted for 89.5% all hits between the PacBio library and the assembly. Of the remaining alignments, 81.3% of hits from X-bin reads were to the X-chromosome assembly and 18% to autosomal arms. From the reads in the A-bin, 94.6% had hits to autosomal arms and only 5.4% to the X-chromosome assembly, with most of these mapping in the 20-25Mbp repeat rich region of the X-assembly, which is known to contain repeats that are shared between the X and the Y chromosome, which drives CQ to autosomal levels (Figure 3) [5]. Since redkmer excludes target sequences that also occur on autosomes or the Y, we next repeated the blast of the X-bin PacBio reads against the genome assembly, after first masking regions of the assembly that have significant similarity to PacBio reads assigned to the A-bin (Table 2). After masking, there remained a significant proportion of hits (27.3%) from reads of the X-bins against the “unknown chromosome”, highlighting that a significant portion of this pseudo-assembly is composed of non-autosomal sequences, or autosomal repeats not represented in the assembly. Of the hits between the X-bin reads, 99% mapped to the X-chromosome assembly and only 0.7% mapped to the autosomal assembly (Table 2). These results indicate that CQ-based prediction of chromosomal origin combined with the exclusion of sequences that overlap with reads assigned to the autosomal bin, can reliably identify X-chromosome specific sequences.

**Table 1:** **Number of hits of PacBio reads to *An. gambiae* genome assembly**

|  | reads | 2R | 2L | 3R | 3L | X | Mt | UN | Y |
| --- | --- | --- | --- | --- | --- | --- | --- | --- | --- |
| A-bin | 1231987 | 426618 | 339163 | 413311 | 276742 | 82397 | 5 | 11747985 | 481 |
| X-bin | 112643 | 3513 | 2107 | 7091 | 2526 | 66134 | 0 | 4710906 | 1 |
| Y-bin | 55541 | 8695 | 1231 | 1716 | 623 | 6970 | 1 | 12378 | 293974 |
| GA-bin | 1300 | 6 | 6 | 12 | 2 | 947 | 0 | 3482 | 0 |
| X-bin <sup>a</sup> | 112643 | 299 | 6 | 2 | 9 | 34248 | 0 | 9438 | 0 |

<sup>a</sup> Xbin after masking genome with reads from A bin

**Figure 3:**
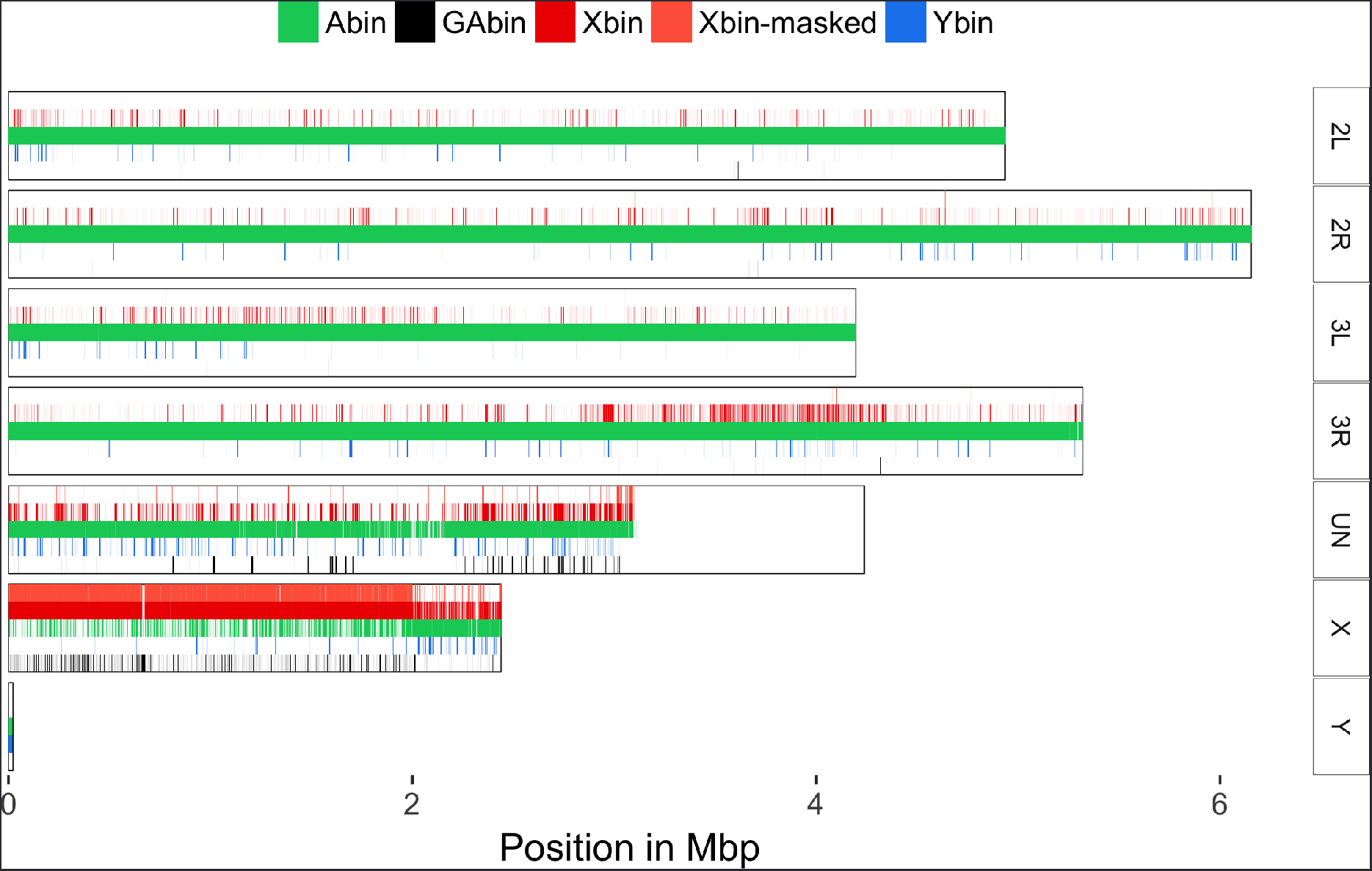
Validation of the long read bin assignment of the Pimperena strain using the AgamP4 genome assembly. Painting of the An. gambiae chromosome assembly with blastn matches longer than 2kbp from reads of each chromosomal bin. The mitochondrial genome is not shown here. The apparent absence of matches between PacBio reads and the “unknown chromosome” downstream of ∼3Mbp results from scaffold length being shorter than the minimum 2kbp alignment length cutoff.

### Candidate X-kmer selection

Redkmer generated 270 million kmers that passed the minimum occurrence threshold in the Illumina data from both males and females (kmernoise=5). The CQ and kmer abundance patterns indicated chromosomal repetitiveness profiles similar to those of the PacBio reads (Figure 4A). Additional redkmer generated plots, providing data on basic statistics of the kmer selection can be found in Supplementary Figures 9-16. Redkmer selected 64619 kmers (0.02%) as being specific to the X-chromosome and within the top 99.5 percentile in kmer abundance (Figure 4B). To identify among these, kmers containing suitable gRNA target sequences for the CRISPR-Cas9 and Cpf1 nucleases we used the FlashFry tool (<u>github.com/aaronmck/FlashFry</u>), using the candidateXkmers.fasta file as input, and all PacBio reads from the A, Y, and GA-bins combined (1288828 reads) as a reference set for performing the CRISPR off-target analysis. 20566 of the candidate X-kmers (32%) were selected as containing suitable gRNA targets for either enzyme, and 669 kmers had sequences suitable for both (Figures 4B and C). Among the 25-mers passing the final selection step, 5 contained the 20bp long gRNA target sequence T1, and 17 contained the 15bp long I-PpoI recognition sequence sites. We have already validated both of these target sequences in independent studies as suitable for inducing sex ratio distortion by X-shredding in *An. gambiae* [3, 4].

**Figure 4:**
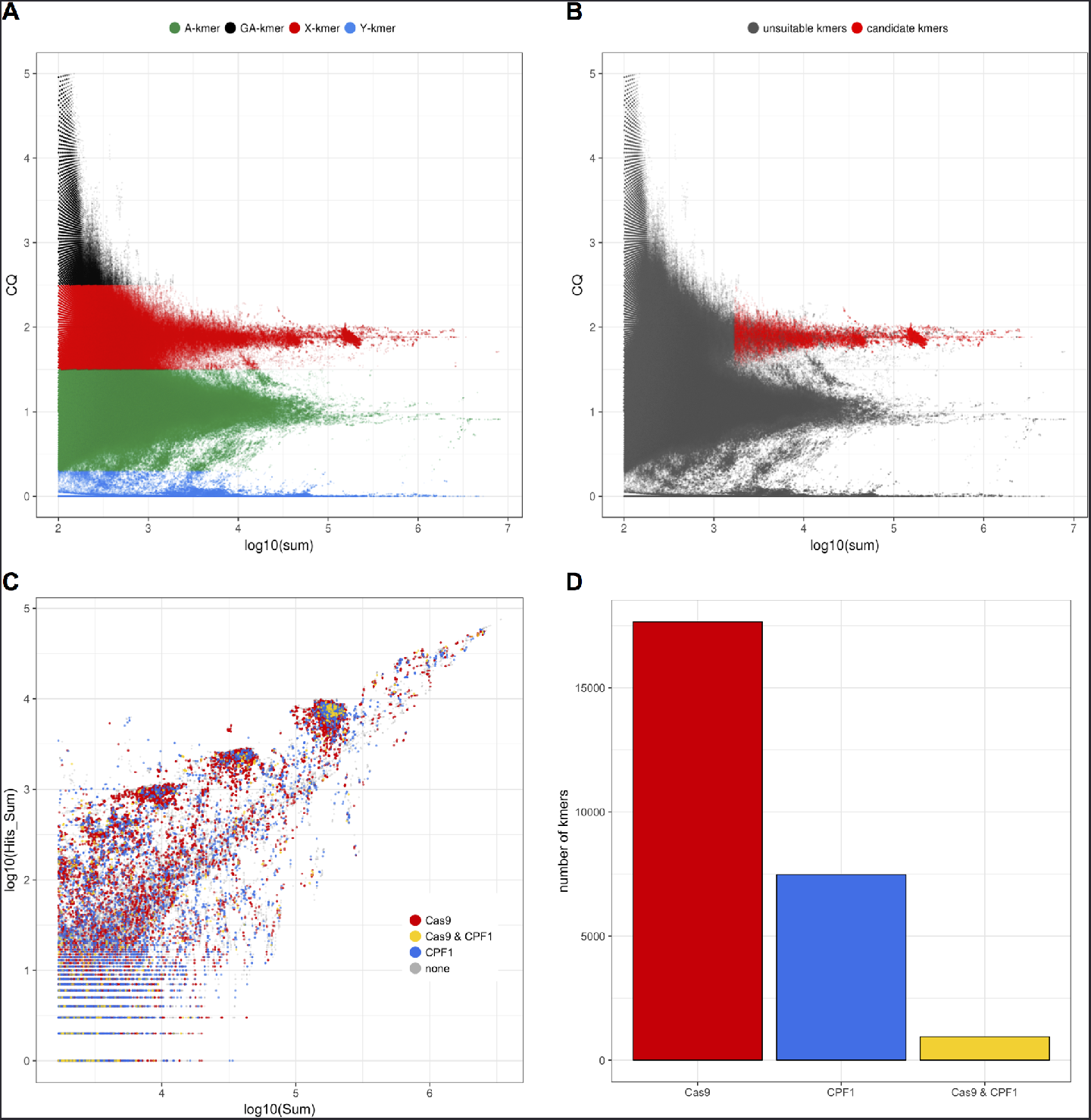
kmer analysis and selection in the Pimperena strain. **A)** kmer-CQ versus abundance (in log10 of sum) in both male and female Illumina libraries for all 270 million redkmer-generated kmers colored by chromosomal bins. **B)** Redkmer selection plot showing kmer-CQ versus abundance for predicted X-chromosome specific and abundant kmers (red dots) versus the unsuitable kmers (grey dots). **C)** Identification of kmers containing sequences predicted to be suitable for targeting by CRISPR endonucleases showing kmer abundance in the Illumina data (x-axis) versus their occurrence within PacBio data (y-axis). Kmers lacking suitable characteristics for CRISPR e.g. the absence of a PAM motif are shown as grey dots, kmers suitable for Cas9 in red, kmers suitable for Cpf1 in blue and those harboring sequences suitable for both Cas9 and Cpf1 are shown in yellow. **D)** Number of kmers containing sequences suitable for CRISPR nuclease platforms.

To evaluate the selection of candidate X-kmers we mapped these using BLAST [1] to each arm of the *An. gambiae* genome assembly (Table 2). Of the 64619 X kmers only 25742 (40%) had hits to the genome assembly, in line with our expectation that many high-copy sequence repeats are poorly represented within genome assemblies. Of the kmers that did have hits within the assembly, 63% mapped to the X chromosome. 78% also matched sequences in the unknown chromosome and 10% had hits to the autosomal arms. Hard filtering of kmers allowing neither perfect nor degenerate off-targets hits to non-X long read bins, reduced the hits to autosomal arms in the assembly to 4.3% (Table 2). Closer inspection of the hits between the X kmers and the autosomes identified two hotspots on chromosome 3R. All 257 kmers mapped exclusively within two genes, AGAP029007 and AGAP029004, both of which represent partial sequences of the 28S ribosomal RNA locus. This is likely an annotation or assembly artifact of the AgamP4 PEST assembly, as we could find no supporting evidence for either gene at the putative positions on chromosome 3R within the Pimperera PacBio reads - none spanned the chromosomal assembly where these two genes are located and none of the kmers mapping to these genes mapped at the junctions where genes meet the flanking assembly.

Based on our use of the *An. gambiae* data set we concluded that redkmer selection, especially after considering the off-target data, is reliable in providing high-confidence target sequences that occur exclusively within the X chromosome. To illustrate this further, we also mapped the 64619 candidate X-kmers against a set of reference sequences using BLAST [1]. As reference sequences we included the entire X-chromosome assembly, the repeats library of *An. gambiae* from Vectorbase and the ribosomal DNA cluster. 61% of these kmers had no hits to any of the reference sequences despite being present within the PacBio reads, highlighting that such sequences are underrepresented within annotated datasets. The ribosomal locus and the X-assembly each accounted for 46% of hits (Figure 5A). With the exception of AgX367, which is a known X-specific satellite in *An. gambiae* [5, 12], all other hits to the reference sequences (22%) were to two families of the site-specific retrotransposons of the R1 and R2 clade that occupy specific positions with 28S rDNA of *An. gambiae* [17, 2, 10] (Figure 5A and B). These results confirm that redkmer is able to identify sequences that are both X-chromosome specific and abundant.

**Table 2:**
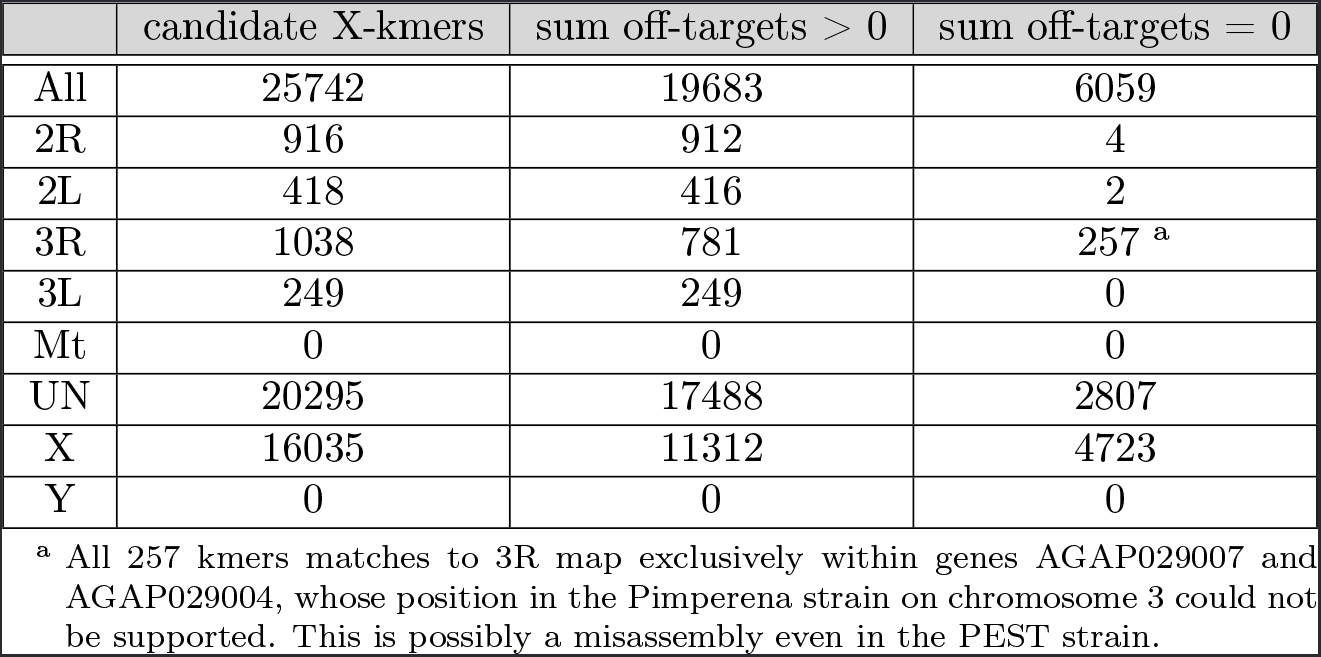
**Number of alignments to the *An. gambiae* PEST assembly**

|  | candidate X-kmers | sum off-targets > 0 | sum off-targets = 0 |
| --- | --- | --- | --- |
| All | 25742 | 19683 | 6059 |
| 2R | 916 | 912 | 4 |
| 2L | 418 | 416 | 2 |
| 3R | 1038 | 781 | 257 <sup>a</sup> |
| 3L | 249 | 249 | 0 |
| Mt | 0 | 0 | 0 |
| UN | 20295 | 17488 | 2807 |
| X | 16035 | 11312 | 4723 |
| Y | 0 | 0 | 0 |

<sup>a</sup> All 257 kmers matches to 3R map exclusively within genes AGAP029007 and AGAP029004, whose position in the Pimperena strain on chromosome 3 could not be supported. This is possibly a misassembly even in the PEST strain.

**Figure 5:**
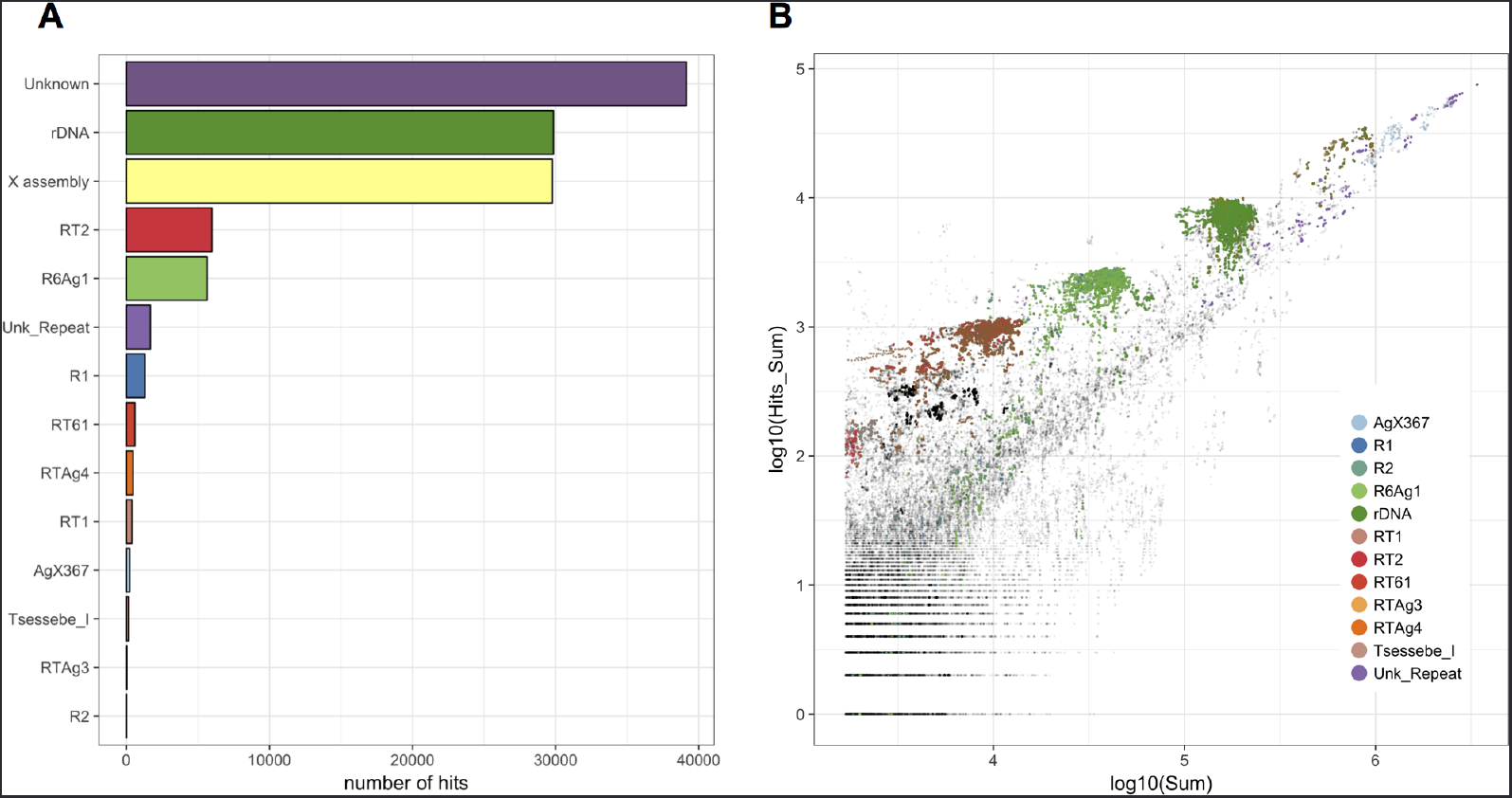
Analysis of the X-candidate kmers of the Pimperena strain. **A)** Blast results showing matches between selected X-kmers and A. gambiae reference sequence collection. **B)** Coverage of selected X-kmers in the Illumina (log10 of Sum) and PacBio data (log10 of Hits_Sum). Each kmer is colored based on the locus it derives from. Because X-kmers corresponding to the X-chromosome assembly do not cluster on the plot, their position is shown but not colored separately.

## Validation of the results using control data from the Asembo1 strain

To test redkmer sensitivity to biological variation we next re-ran the pipeline using Illumina WGS libraries from males and female of the Asembo1 strain of *An. gambiae*. Colonized in 1997 in Asembo, Kenya the rDNA is believed to have introgressed onto the Y chromosome in this strain forming a Mopti/Savanna hybrid in males (Wilkins et al 2007). Cytological evidence confirms that this strain harbors the ribosomal gene cluster not only on the X-chromosome but also on the Y (Hall et al 2016). Long read single-molecule sequencing is not available for the Asembo1 strain, so we evaluated its Illumina data against the Pimperena PacBio data set. We expected that CQ based prediction of chromosomal origin would result in significantly different chromosome bins, particularly for reads corresponding to the rDNA cluster and its associated repeats. The highly repetitive cloud of PacBio reads previously assigned to the X bin using Pimperena Illumina data (Figure 2B) was absent when mapping was done using the Asembo1 reads (Figure 6A). Effectively, PacBio reads corresponding to the X-heterochromatic region, were now being assigned to the autosome bin, as co-occurrence of sequences on the X and Y chromosome drives CQ to autosomal levels. Interestingly, the repeat content and abundance Y-chromosome assigned reads did not indicate that the X-Y recombination event that transferred the ribosomal cluster from the X to the Y was reciprocal (Figure 6A). Redkmer now identified only 210 kmers as specific to the X chromosome (Figure 6B). We found by blasting these kmers to the references sequences used above, that 120 of these kmers (57%) had no hits. Of the remaining 90 kmers, 74 had hits to the X-chromosome assembly, 89 matched the RT2 or R7Ag1 retroposons that insert specifically within the rDNA locus, and 16 had hits to the rDNA cluster. The X and Y chromosomal rDNA loci of Asembo1 are known to be polymorphic and contain mixed arrays of both *An. gambiae* and An. coluzzi (called M and S in that study) [21]. We reasoned that redkmer did not exclude these 94 kmers because these occurred specifically within the X-chromosome array and not on the Y chromosome rDNA array. To confirm this, we selected a subset of all kmers matching to the rDNA reference sequence (611 of 261743585 kmers) and found that the majority did indeed display kmer CQ indicative of linkage to both sex-chromosomes, but that a small number did retain CQ values higher than 1.5 (the X-CQ cutoff) (Figure 6B).

**Figure 6:**
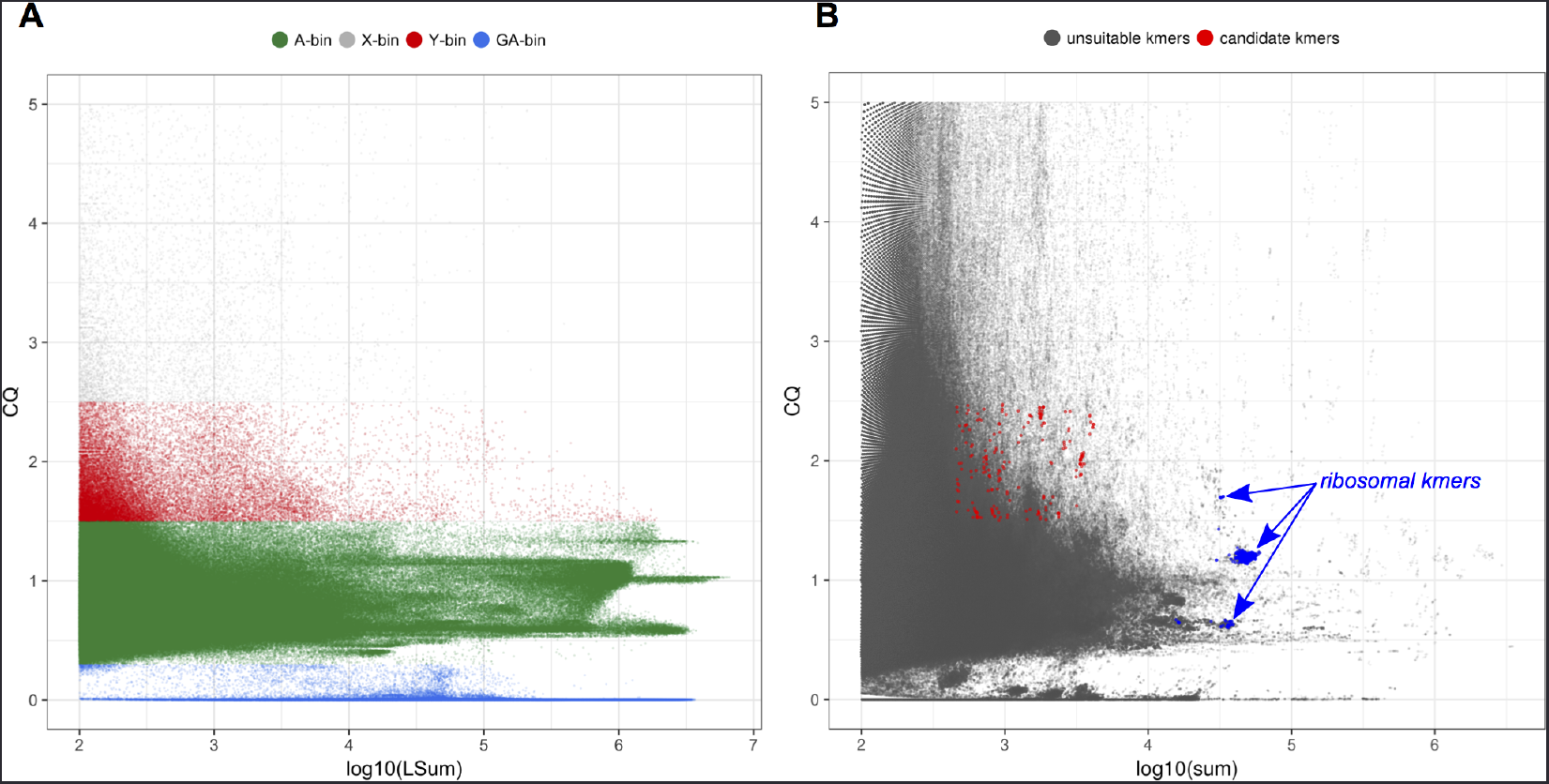
Analysis of the Asembo1 strain. **A)** PacBio read CQ (ratio of female/male data) over log10 of LSum (total length-normalized number of mapping reads from male and female illumina data) showing absence of X-specific repeat cloud in the Asembo strain. B) Plot showing kmer-CQ versus abundance for selected X-chromosome specific and abundant kmers (red dots - candidate kmers) and those not suitable passing selection(grey dots). Blue dots indicate kmers mapping to the ribosomal array confirming that this locus is not X-specific in the Asembo1 strain (CQ ~ autosomal) and thus unsuitable for X-shredding. The 89 kmers that are selected and map to this locus have also been highlighted (top blue arrow).

## Discussion

Knowledge of X-chromosome specific sequences is an essential component for the development of synthetic sex ratio distorters based on X-chromosome shredding. For the vast majority of species this type of information is not available however, owing to the difficulty of characterizing repetitive, heterochromatic DNA, whose characteristics make genome assembly and scaffolding unreliable. The advent of next-generation sequencing (NGS) and more recently single-molecule sequencing is beginning to provide the technologies required to study the makeup and properties of heterochromatic sequences. For example, we have previously shown that combining long single-molecule data with short Illumina sequencing can be a successful strategy to characterize both the genic and repetitive content of the Y chromosome of mosquitoes [5], and similar methods are now being applied to other species. However, an approach to tackle X chromosome specific repetitive sequences, which is complicated by the fact that X chromosomes sequences unlike those on the Y chromosome are not sex specific, has not been developed. To address this gap we developed the redkmer pipeline which is designed to require as input only raw WGS data with minimal filtering and error-correction. The main output is a tab-delimited file providing data on kmers selected as X-chromosome specific and abundant, including kmer-CQ, coverage, X-chromosome-specificity and off-targeting data, along with a fasta file for downstream analysis. In addition, redkmer data can be used to infer and identify X and Y chromosome sequences within the “chromosomal bins” of PacBio reads. Plots describing the PacBio reads and the kmers are also produced at the end of the redkmer pipeline (Supplementary Fig. 1-16). While neither the assignment of Pacbio reads to chromosomal bins nor the calculation of kmer CQ values are fully reliable on their own, we show that the combination of these two strategies can be used to efficiently and reliably identify X-chromosome specific kmer sequences.

The *An. gambiae* Pimperena Illumina dataset consists of ∼140M 100bp Illumina reads per sex and ∼2M Pacbio reads that passed length filtering. The combination of these reads results in >10^14^ possible cross alignments, which is why redkmer is implemented for parallel execution by splitting of the Pacbio input data. To test the limits of the pipeline we have also tested redkmer on another data set featuring ∼400M Illumina reads per sex and unpublished ∼8M Pacbio reads (data not shown) and confirmed that the pipeline is able to handle the larger datasets that are now becoming available. We showed that running redkmer with data from *An. gambiae* correctly identified known X-chromosome specific and abundant sequences, some of which we used previously to develop the first synthetic sex ratio distorters by X-chromosome shredding Galizi et al. [3]. We also showed using the control strain Asembo1, in which a large fraction of X-chromosome repetitive sequences are shared with the Y, that redkmer target prediction differs in line with our expectations. Therefore running redkmer with data from different species in the future may help identify those that are suitable for the development of X-shredder based genetic control strategies, and identify target sites for doing so.

## Conclusion

The data presented in this study have shown that combination of single-molecule and long-read sequencing when combined with short read WGS data from males and females can be used to efficiently and reliably identify X-chromosome specific sequences, which can be used to develop X-shredder based sex ratio distortion systems.

## Acknowledgements

We would like to thank Matt J Harvey and Adam Phillippy for helpful suggestions in building the redkmer pipeline. P.A.P. was supported by a Rita Levi Montalcini award from the Ministry Education, University and Research (MIUR – D.M. no. 79 04.02.2014). Funded by the BBSRC under the research grant BB/P000843/1 to N.W. Data used in this study were funded in part by a grant from the Foundation for the National Institutes of Health through the Vector-Based Control of Transmission: Discovery Research (VCTR) program of the Grand Challenges in Global Health initiative of the Bill & Melinda Gates Foundation. This study was funded by the European Research Council under the European Union’s Seventh Framework Programme ERC grant no. 335724 awarded to N.W.

## References

[1] Altschul S. F., Gish W., Miller W., Myers E. W. and Lipman D. J. [1990], ‘Basic local alignment search tool’, J Mol Biol 215(3), 403–10.

[2] Besansky N. J., Paskewitz S. M., Hamm D. M. and Collins F. H. [1992], ‘Distinct families of site-specific retrotransposons occupy identical positions in the rrna genes of anopheles gambiae’, Mol Cell Biol 12(11), 5102–10.

[3] Galizi R., Doyle L. A., Menichelli M., Bernardini F., Deredec A., Burt A., Stoddard B. L., Windbichler N. and Crisanti A. [2014], ‘A synthetic sex ratio distortion system for the control of the human malaria mosquito’, Nat Commun 5, 3977.

[4] Galizi R., Hammond A., Kyrou K., Taxiarchi C., Bernardini F., O’Loughlin S. M., Papathanos P.-A., Nolan T., Windbichler N. and Crisanti A. [2016], ‘A crispr-cas9 sex-ratio distortion system for genetic control’, Sci Rep 6, 31139.

[5] Hall A. B., Papathanos P.-A., Sharma A., Cheng C., Akbari O. S., As-sour L., Bergman N. H., Cagnetti A., Crisanti A., Dottorini T., Fioren-tini E., Galizi R., Hnath J., Jiang X., Koren S., Nolan T., Radune D., Sharakhova M. V., Steele A., Timoshevskiy V. A., Windbichler N., Zhang S., Hahn M. W., Phillippy A. M., Emrich S. J., Sharakhov I. V., Tu Z. J. and Besansky N. J. [2016], ‘Radical remodeling of the y chromosome in a recent radiation of malaria mosquitoes’, Proc Natl Acad Sci U S A 113(15), E2114–23.

[6] Hall A. B., Qi Y., Timoshevskiy V., Sharakhova M. V., Sharakhov I. V. and Tu Z. [2013], ‘Six novel y chromosome genes in anopheles mosquitoes discovered by independently sequencing males and females’, BMC Genomics 14, 273.

[7] Hamilton W. D. [1967], ‘Extraordinary sex ratios. a sex-ratio theory for sex linkage and inbreeding has new implications in cytogenetics and entomology’, Science 156(3774), 477–88.

[8] Hammond A. M., Kyrou K., Bruttini M., North A., Galizi R., Karls-son X., Carpi F., D’Aurizio R., Crisanti A. and Nolan T. [2017], ‘The creation and selection of mutations resistant to a gene drive over multiple generations in the malaria mosquito’, bioRxiv. URL: http://www.biorxiv.org/content/early/2017/06/12/149005.1.

[9] Holt R. A., Subramanian G. M., Halpern A., Sutton G. G., Charlab R., Nusskern D. R., Wincker P., Clark A. G., Ribeiro J. M. C., Wides R., Salzberg S. L., Loftus B., Yandell M., Majoros W. H., Rusch D. B., Lai Z., Kraft C. L., Abril J. F., Anthouard V., Arensburger P., Atkinson P. W., Baden H., de Berardinis V., Baldwin D., Benes V., Biedler J., Blass C., Bolanos R., Boscus D., Barnstead M., Cai S., Center A., Chaturverdi K., Christophides G. K., Chrystal M. A., Clamp M., Cravchik A., Curwen V., Dana A., Delcher A., Dew I., Evans C. A., Flanigan M., Grundschober-Freimoser A., Friedli L., Gu Z., Guan P., Guigo R., Hillenmeyer M. E., Hladun S. L., Hogan J. R., Hong Y. S., Hoover J., Jaillon O., Ke Z., Kodira C., Kokoza E., Koutsos A., Letu-nic I., Levitsky A., Liang Y., Lin J.-J., Lobo N. F., Lopez J. R., Malek J. A., McIntosh T. C., Meister S., Miller J., Mobarry C., Mongin E., Murphy S. D., O’Brochta D. A., Pfannkoch C., Qi R., Regier M. A., Remington K., Shao H., Sharakhova M. V., Sitter C. D., Shetty J., Smith T. J., Strong R., Sun J., Thomasova D., Ton L. Q., Topalis P., Tu Z., Unger M. F., Walenz B., Wang A., Wang J., Wang M., Wang X., Woodford K. J., Wortman J. R., Wu M., Yao A., Zdobnov E. M., Zhang H., Zhao Q., Zhao S., Zhu S. C., Zhimulev I., Coluzzi M., della Torre A., Roth C. W., Louis C., Kalush F., Mural R. J., Myers E. W., Adams M. D., Smith H. O., Broder S., Gardner M. J., Fraser C. M., Birney E., Bork P., Brey P. T., Venter J. C., Weissenbach J., Kafatos F. C., Collins F. H. and Hoffman S. L. [2002], ‘The genome sequence of the malaria mosquito anopheles gambiae’, Science 298(5591), 129–49.

[10] Kojima K. K. and Fujiwara H. [2003], ‘Evolution of target specificity in r1 clade non-ltr retrotransposons’, Mol Biol Evol 20(3), 351–61.

[11] Koren S., Walenz B. P., Berlin K., Miller J. R., Bergman N. H. and Phillippy A. M. [2017], ‘Canu: scalable and accurate long-read assembly via adaptive k-mer weighting and repeat separation’, Genome Res 27(5), 722–736.

[12] Krzywinski J., Sangaré D. and Besansky N. J. [2005], ‘Satellite dna from the y chromosome of the malaria vector anopheles gambiae’, Genetics 169(1), 185–96.

[13] Langmead B. and Salzberg S. L. [2012], ‘Fast gapped-read alignment with bowtie 2’, Nat Methods 9(4), 357–9.

[14] Langmead B., Trapnell C., Pop M. and Salzberg S. L. [2009], ‘Ultra-fast and memory-efficient alignment of short dna sequences to the human genome’, Genome Biol 10(3), R25.

[15] Li H., Handsaker B., Wysoker A., Fennell T., Ruan J., Homer N., Marth G., Abecasis G., Durbin R. and 1000 Genome Project Data Processing Subgroup [2009], ‘The sequence alignment/map format and samtools’, Bioinformatics 25(16), 2078–9.

[16] Marçais G. and Kingsford C. [2011], ‘A fast, lock-free approach for efficient parallel counting of occurrences of k-mers’, Bioinformatics 27(6), 764–70.

[17] Paskewitz S. M. and Collins F. H. [1989], ‘Site-specific ribosomal dna insertion elements in anopheles gambiae and a. arabiensis: nucleotide sequence of gene-element boundaries’, Nucleic Acids Res 17(20), 8125–33.

[18] S, A. [n.d.], Fastqc: a quality control tool for high throughput sequence data andrewss http://www.bioinformatics.babraham.ac.uk/projects/fastqc/ added by an anonymous collaboratoredit / delete open in paperpile fastqc: a quality control tool for high throughput sequence data. http://www.bioinformatics.babraham.ac.uk/projects/fastqc/.

[19] Schliekelman P., Ellner S. and Gould F. [2005], ‘Pest control by genetic manipulation of sex ratio’, J Econ Entomol 98(1), 18–34.

[20] Unckless R. L., Clark A. G. and Messer P. W. [2017], ‘Evolution of resistance against crispr/cas9 gene drive’, Genetics 205(2), 827–841.

[21] Wilkins E. E., Howell P. I. and Benedict M. Q. [2007], ‘X and y chromosome inheritance and mixtures of rdna intergenic spacer regions in anopheles gambiae’, Insect Mol Biol 16(6), 735–41.

